# Proteomic analysis reveals different sets of proteins expressed during high temperature stress in two thermotolerant isolates of *Trichoderma*

**DOI:** 10.1101/2021.08.12.456037

**Authors:** Sowmya Poosapati, Viswanathaswamy Dinesh Kumar, Ravulapalli Durga Prasad, Monica Kannan

## Abstract

Several species of the soil borne fungus of the genus *Trichoderma* are known to be versatile, opportunistic plant symbionts, and are the most successful biocontrol agents used in today’s agriculture. To be successful in the field conditions, the fungus must endure varying climatic conditions. Studies have indicated that high atmospheric temperature coupled with low humidity is a major limitation for the inconsistent performance of *Trichoderma* under field conditions. Understanding the molecular modulation associated with such *Trichoderma* that persist and deliver under abiotic stress condition will aid in exploiting the worth of these organisms for such use. In this study, comparative proteomic analysis using two-dimensional gel electrophoresis (2DE) and matrix-assisted laser desorption/time-of-flight (MALDI-TOF-TOF) mass spectrometry was used to identify proteins associated with thermotolerance in two thermotolerant isolates of *Trichoderma*—*T. longibrachiatum* 673, TaDOR673 and *T. asperellum* 7316, TaDOR7316—and 32 differentially expressed proteins were identified. Sequence homology and conserved domains were used to identify these proteins and to assign probable function to them. Thermotolerant isolate, TaDOR673, seemed to employ the stress signaling MAPK pathways and heat shock response pathways to combat the stress condition whereas the moderately tolerant isolate, TaDOR7316, seemed to adapt to high temperature conditions by reducing the accumulation of mis-folded proteins through unfolded protein response pathway and autophagy. Also, there were unique as well as common proteins that were differentially expressed in the two isolates studied.

## 1. Introduction

Fungi belonging to the genus *Trichoderma* account for more than 60% of all the registered biopesticides (Verma et al. 2007). *Trichoderma* is used to antagonize phytopathogenic fungi and the antifungal properties of *Trichoderma* are principally due to their ability to produce antibiotics (Vinale et al. 2008) and/or hydrolytic enzymes (Benitez et al. 2004) and competition for space and nutrients (Elad 2000). Even if *Trichoderma* species can promote plant growth and induce resistance to biotic and abiotic stresses (Hermosa et al. 2012), their role as bio-pesticide has primarily contributed to their commercial success as bio-agents. Nevertheless, *Trichoderma* by themselves are not immune to abiotic stress like moisture deficiency, higher temperature, etc., that tend to cause morphological, physiological, biochemical and molecular changes and adversely affect the beneficial consequences of these bioagents. For example, soil hydrological conditions influence the growth and antagonistic properties of *Trichoderma* (Tronsmo and Dennis 1978; Luard and Griffin 1981; Magan 1988) and soil temperature affect the radial extension of *Trichoderma* (Knudsen and Bin 1990).

Further, agricultural practices such as solarisation of soil have been widely used to eradicate infectious pathogens from soil, (Katan 1981) as these practices weaken the potential of pathogens to damage crops and increase their susceptibility to bio-agents. Such practices also subject *Trichoderma*, if applied to the soil, to high temperature stress. Therefore, there is a synergistic benefit in combining sub-lethal solarisation with thermotolerant bioagents, especially *Trichoderma*, to suppress crop-damaging, temperature-tolerant pathogens of soil (Tjamos and Fravel, 1995; Tjamos and Paplomatas, 1987).

Thermotolerant strains have been identified in *Bacillus* and *Pseudomonas* (Kumar et al. 2014), but as *Trichoderma* is already a widely used bio-agent, identification of thermotolerant strains of this genus will be relevant apart from the economic significance entailed (Poosapati et al. 2014). Advancements in ‘omics’ sciences have enabled researchers to understand the mechanisms of thermotolerance in organisms such as *Saccharomyces cerevisiae*, *Metarhizium anisopliae* etc. (Shui W, 2015; Wang et al. 2014; Verghese J, 2012; Tereshina VM, 2005) and this information has been exploited to improve the strains and stress resistance in plants (Nicolas et al. 2014; Montero-Barrientos et al. 2007 & 2008). In *Trichoderma*, specifically, the *hog* 1 gene (ThHog1) from MAPK pathway was the first to be implicated in tolerance to osmotic stress (Delgado-Jarana et al. 2006). Molecular studies have revealed that components of cell wall undergo a dynamic change to adapt to stress conditions (Ram et al. 2004; Chatterjee et al. 2000) and act as mediators of stress response (Konopka 2012; Chen et al. 2008; Yang et al. 2003) to activate the cell wall integrity (CWI) MAPK pathway (Chauhan & Calderone 2008; Chauhan et al. 2006). Specifically, Protein kinase C (Pkc) is implicated in the maintenance of cell wall integrity (CWI) in response to different environmental stresses (Colabardini et al. 2014).

Fungi are known to develop different strategies involving diverse regulatory mechanisms to adapt to stress conditions. Heat shock response is a highly conserved pathway that results in the immediate synthesis of a pool of cytoprotective genes in the presence of diverse environmental stresses. Accumulation of misfolded proteins rapidly activates Hsf1 (Verghese et al. 2012; Abrams et al. 2014; Gowda et al. 2013) which in turn induces several heat shock proteins that help in proper folding of misfolded proteins during the stress conditions (Kim et al. 2007; Yin et al. 2009). Role of Hsf1 in cell wall remodeling (Imazu and Sakurai, 2005) had indicated a cross talk between stress induced pathways. Only a few researchers have evaluated the role of *Trichoderma* derived heat shock proteins in tolerance to heat, salt and oxidative stress (Montero-Barrientos et al. 2007 and 2008). Other genes like glutathione transferase (Dixit et al. 2011) and proteins with glucosidase activity (Hermosa et al. 2011) were also shown to enhance tolerance to several abiotic stresses. Studies were highly focused to understand the effect of various stresses in pathogenic fungi of clinical importance (Yin et al. 2009; Schmidt et al. 2008; Kusch et al. 2007; Liu et al. 2014). Nevertheless, researchers have reviewed the role of various *Trichoderma* genes in plant stress tolerance (Nicolas et al. 2014).

In spite of the growing genomic resources in *Trichoderma* species (genomes of several *Trichoderma* species have been sequenced [http://genome.jgi-psf.org/Triat1/Triat1.home.html]) and ESTs from several species of *Trichoderma* available in TrichoEST database (Vizcaíno et al. 2006; Rey et al. 2004), much of the research so far is focused on studies involving plants-pathogen-bioagent interactions (Marra et al. 2006), mycoparasitim (Druzhinina et al. 2011), biocontrol related genes and enzymes (Srivastava et al. 2014; Sharma et al. 2011) and proteases produced by bio-agents (Atanasova et al. 2013; Rawlings et al. 2012; Suarez et al. 2007; Benítez et al. 2004). But there are few or no reports on molecular changes associated with heat stress conditions in *Trichoderma*.

In the present investigation, we used two thermotolerant *Trichoderma* strains identified previously in our lab (Poosapati et al. 2014) that differed in their level of tolerance to temperature stress. Proteins that were differentially expressed, when subjected to temperature stress, were identified using 2D electrophoresis and MALDI-TOF. To the best of our knowledge, this is the first report of proteomic analysis of heat stress tolerance in *Trichoderma* species.

## 2. Materials and methods

### 2.1. Strains

The thermotolerant strains of *Trichoderma viz*., *T. longibrachiatum* 673, TaDOR673 and *T. asperellum* 7316, TaDOR7316, which were identified previously and stored in Microbial type culture collection (MTCC) (Poosapati et al. 2014), were used for this study. These thermotolerant strains were identified from a pool of *Trichoderma* isolates, isolated from soil samples collected from various regions of India. To confirm their identity, multi-locus sequencing (elongation factor1 alpha and RNA polymerase subunit B) was performed and the results confirmed the identity of these strains (Accession numbers provided in supplementary table 1).

### 2.2. Growth conditions

The strains were grown on potato dextrose agar at 28 °C for 7 days. Briefly, 2-3 mycelial discs from 7-day-old culture were used to inoculate 100 ml of potato dextrose broth in 250 ml Erlenmeyer flasks and were incubated for 3 days at 28 °C and 200 rpm followed by incubation in a static position for 4 days to allow better sporulation. Two biological replicates were used and the cultures were exposed to a temperature of 48 °C for 1 h and 4 h (treated samples). After incubation, mycelium was filtered and frozen in liquid nitrogen and stored at −80 °C, until use. Samples from both the strains that were not subjected to thermal stress were used as control.

### 2.3. Protein extraction

Protein extraction protocol described by Jacobs et al. (2001) was followed, with a few modifications. In brief, approximately 1 g of fungal mycelium was ground in liquid nitrogen and suspended in 10 ml of cold (−20 °C) acetone solution containing 13.3 % Trichloroacetic acid (TCA) and 0.07 % ß-mercaptoethanol. Samples were vortexed and maintained at −20 °C for at least 3 h with intermittent shaking to allow the precipitation of protein and then centrifuged at 14000 rpm for 20 min at 4 °C. The pellets were washed three times with cold (−20 °C) acetone solution containing 0.07 % ß-mercaptoethanol and air dried. The pellets were resuspended in labeling buffer (8 M urea, 2 M thiourea, 2 % 3-[(3-cholamidopropyl)-dimethyl-ammonio]-1-propane sulfonate (CHAPS) and 1 % DTT, mixed and kept on an orbital shaker for 2 h at 37 °C to obtain complete protein solubilization. Samples were centrifuged (14000 rpm, 60 min at 20 °C) and the supernatants were recovered. The supernatant was desalted using PD spin trap columns (GE Healthcare, USA), stored at −80 °C until use and protein concentration was determined by Bradford protein assay using bovine serum albumin (BSA) as a standard.

### 2.4. Two-dimensional electrophoresis (2DE) and image analysis

Isoelectric focusing (IEF) was performed in 18 cm immobilized-pH-gradient (IPG) strips (GE Healthcare, USA) with a pH range of 4-7 L, rehydrated in a solution of 7 M Urea, 2 M thiourea, 4 % CHAPS, 1 % DTT, 2 % carrier ampholytes and 1 x Protease inhibitor cocktail (Sigma Aldrich, USA). About 100 μl of total protein solution (equivalent to 500 μg) was loaded onto the focusing tray and was let to be absorbed into the gel strip. The IPG strips were focused upto a total of 10 KVh using a five step program (step 500 V-for 3 h, gradient-500 V for 5 h, gradient 10 KV for 8 h and 60 KVh was reached, then finally 500 V for 10 h) in Ettan^™^ IPGphor^™^ isoelectric focusing system (Amersham Biosciences, USA). Focused strips were equilibrated by placing them in a solution of 6 M urea, 0.05 M Tris-HCl, pH 8.8, 20 % glycerol, 2 % Sodium dodecyl sulphate (SDS), 2 % Dithiothreitol (DTT) for 10 min and then in 6 M urea, 0.05 M Tris-HCl, pH 8.8, 20 % glycerol, 2 % SDS, 2.5 % iodo-acetamide for 10 min more. For the second dimension SDS-PAGE, the IPG strips were loaded on top of 12.5 % polyacrylamide gel in Ettan^™^ DALTsix Large vertical system (Amersham Biosciences, USA). Polyacrylamide gels were then electrophoresed at a constant voltage of 150 V for 60 min in Tris-glycine-SDS buffer, fixed in 40: 10 % v/v methanol: acetic acid (overnight) and stained with sensitive colloidal Coomassie blue G-250. Gels were destained with water until the background was clear and stored in 40: 10 % methanol: acetic acid solution until further use. Protein patterns in the polyacrylamide gels were recorded as digitized images using a calibrated densitometric scanner (GE Healthcare, USA) and analyzed (normalization, spot matching, expression analyses, and statistics) using Image Master 2-D Platinum 6 image analysis software (GE Healthcare, USA). Spots on the gel were identified and each spot was assigned a spot quantity (q), an approximate amount of protein based on relative spot size and intensity. Differential expression (DE) was measured as the relative ratio of q for the same spot between two comparative gels.

### 2.5. Mass spectrometry and protein identification

In-gel digestion and matrix-assisted laser desorption/ionization time of flight mass spectrometric (MALDI-TOF MS) analysis was done with a MALDI-TOF/TOF mass spectrometer (Bruker Autoflex III smartbeam, Bruker Daltonics, Bremen, Germany) according to the method described by Shevchenko et al. (1996) with slight modifications. Two technical replicates (for each of the two biological replicates) representing each treatment were used and the differentially expressed spots in the comparative gels were identified and manually excised from the gels. The excised gel pieces were then destained with 100 μl of 50 % acetonitrile (ACN) in 25 mM ammonium bicarbonate (NH_4_HCO_3_) at least five times. Thereafter, the gel pieces were treated with 10 mM DTT in 50 mM NH_4_HCO_3_ and incubated at 56 °C for an hour. This was followed by treatment with 55 mM iodo-acetamide in 50 mM NH_4_HCO_3_ for 45 min at room temperature in dark (25 ± 2 °C), washed with 25 mM NH_4_HCO_3_ and ACN, dried in Speed Vac and rehydrated in 20 μl of 25 mM NH_4_HCO_3_ solution containing 12.5 ng/μl trypsin (sequencing grade, Promega, Wisconsin, USA). This mixture was incubated on ice for 10 min and digested at 37 °C overnight. After digestion, the mixture was pulse spun for 10 s and the supernatant collected in a fresh Eppendorf tube. The gel pieces were re-extracted with 50 μl of 0.5 % trifluoroacetic acid (TFA) and ACN (1: 1) for 15 min with frequent mixing. The supernatants were then pooled together and dried using SpeedVac and were reconstituted in 5 μl of 1: 1 ACN and 0.1 % TFA. About 2 μl of the above sample was mixed with 2 μl of freshly prepared a-cyano-4-hydroxycinnamic acid (CHCA) matrix in 50 % ACN and 0.1 % TFA (1: 1) and 1 μl was spotted onto the target plate.

Proteins were identified through database searches (PMF and MS/MS) using MASCOT program (http://www.matrixscience.com) employing Biotools software version 3.0 (Bruker Daltonics). Homology search using mass values was done with existing digests and sequence information from NCBInr and Swiss-Prot database by setting the taxonomic category to “Other fungi” and setting the search parameters to fixed modification of carbamidomethyl (C), variable modification of oxidation (M), enzyme trypsin, peptide charge of 1 ^+^ and monoisotropic. According to the MASCOT probability analysis (p < 0.05), only significant ‘hits’ were accepted for protein identification.

For proteomic analysis, spots were analyzed using Image Master 2-D Platinum image analysis software (GE Healthcare). One-way factor ANOVA (P < 0.05) was performed by keeping the values of the best-matched replicate gels from the two independent experiments. Normalized volume (% vol) of each spot was automatically calculated by the Image analysis software.

## 3. Results and Discussion

### 3.1. Morphological differences of thermotolerant isolates

In the present investigation, two thermotolerant isolates of *Trichoderma*—*T. longibrachiatum* 673 (TaDOR673) and *T. asperellum* 7316 (TaDOR7316)—that were identified earlier at IIOR, were used. These isolates were able to tolerate a heat shock of 52 °C for 4 h and were able to retain their morphological features after recovery from heat stress. The level of thermotolerance, however, was distinct between these isolates. The isolate TaDOR673 was highly tolerant to the heat shock condition tested, with a mean spore count (log c.f.u/ml) of 4.33 after the treatment whereas TaDOR7316 had a mean spore count of 1.16. The two isolates exhibited distinct morphology. TaDOR673 was a sparsely sporulating fungus with yellow pigmentation whereas TaDOR7316 was a dense sporulating fungus (Fig 1). TaDOR673 was able to tolerate higher thermal stress even at hyphal stage when compared to TaDOR7316 (Poosapati et al. 2014)

**Fig. 1.**
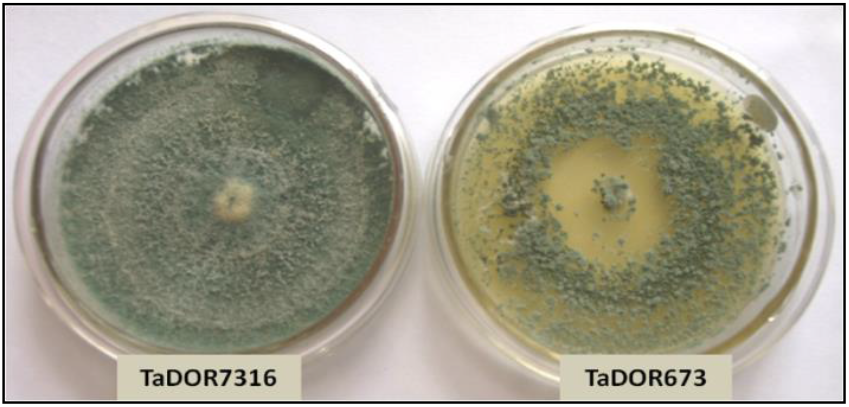
Morphology of thermotolerant isolates of *Trichoderma*

Biochemical analysis revealed that both these isolates accumulate a higher concentration of trehalose, a known compatible solute, during heat shock (Poosapati et al. 2014). We hypothesized that these morphological and physiological differences might contribute to their distinct levels of thermotolerance and we used proteomic approaches to discern the changes associated with thermal stress in these two isolates of *Trichoderma*.

### 3.2. Protein profiling of thermotolerant isolates

At temperatures higher than 48 °C significant reduction in protein abundance was observed (data not shown) in both the isolates and hence incubation temperature of 48 °C was selected to perform the heat stress studies and total protein was isolated from the strains exposed to 48 °C after 1h and 4h. We appreciate that some transiently up-regulated proteins, either within one hour of exposure or those which are altered between 1 h and 4 h, might have been missed from our analyses, as has been emphasised by Kusch et al. (2007)

Using coomassie brilliant blue staining, approximately 580 proteins were visualized in 2DE gels from each isolate (Fig 2, 3). Two replicate gels were used to construct a representative master gel (RMG) for each sample. Protein spots with significant changes in expression-levels were considered to be important for thermotolerance (Fig 4 & Supplementary Fig 1). A total of 32 protein spots were identified from both the isolates using MALDI-TOF-TOF (Table 1, 2). Among those proteins, proteins related to heat shock response were found to be commonly affected in both the isolates. As most of the protein hits were obtained with other fungi, search against *Trichoderma* was also carried out using BLASTP to identify the homologous proteins (Supplementary table 2, 3). The presence of highly conserved domains was taken into consideration using NCBI’s Conserved Domain Database (CDD).

**Fig. 2.**
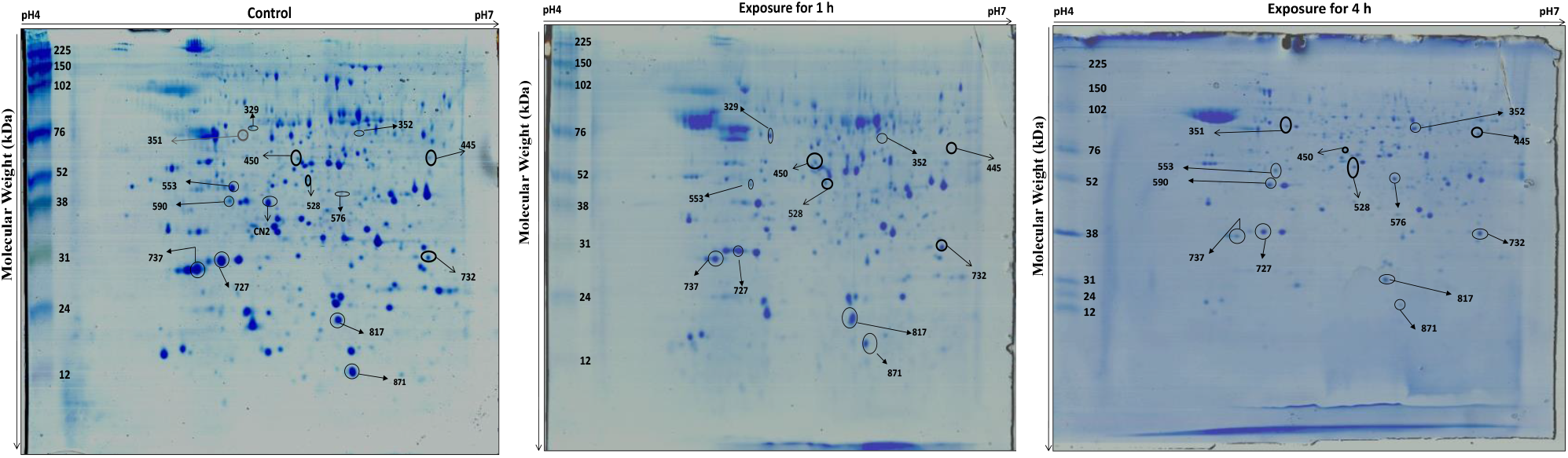
Two-dimensional gel image of the protein expression pattern in *T. longibrachiatum* 673, TaDOR673 subjected to thermal stress. The gels were obtained in duplicates; a representative of each duplicate is shown. Identified protein spots are numbered and listed in Table 1. The pH gradient is marked above the gel and the molecular mass protein standards (kDa) are indicated on the left of the gel

**Fig. 3.**
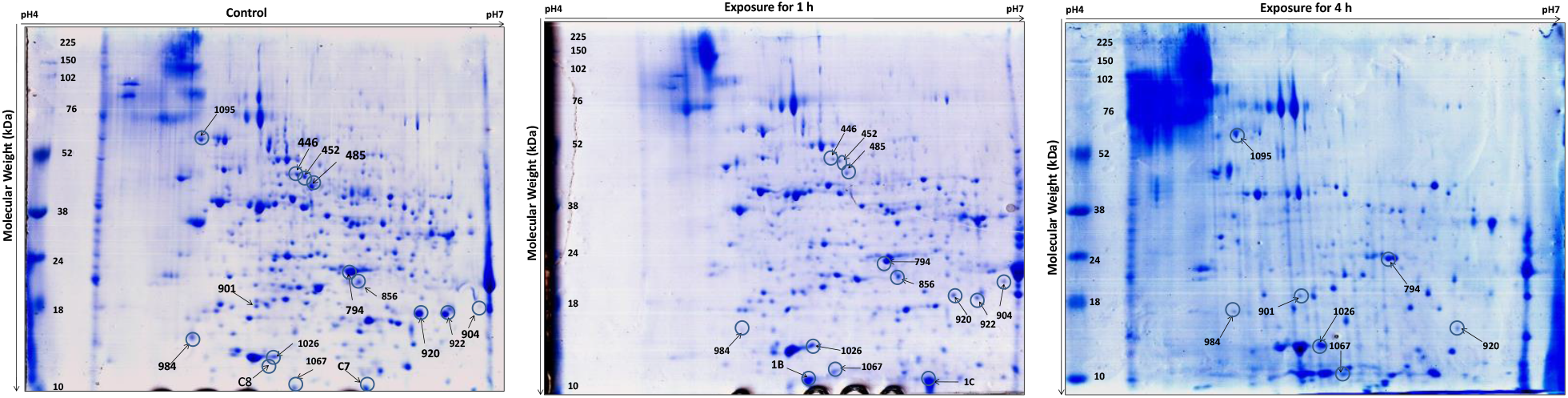
Two-dimensional gel image of the protein expression pattern in *T. asperellum* 7316, TaDOR7316 subjected to thermal stress. The gels were obtained in duplicates; a representative of each duplicate is shown. Identified protein spots are numbered and listed in Table 2. The pH gradient is marked above the gel and the molecular mass protein standards (kDa) are indicated on the left of the gel

**Fig. 4.**
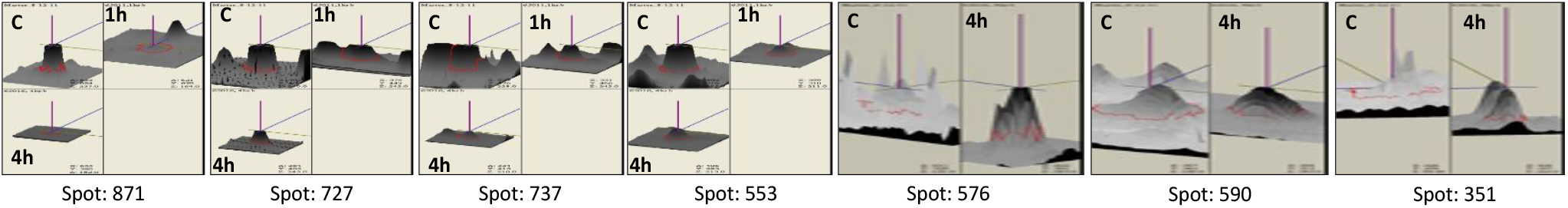
3D view of differential intensity levels of some of the protein spots identified in thermotolerant isolates of *Trichoderma* during the heat stress. Spot intensity was quantified by Image Master 2-D Platinum V6.0 image analysis software (GE Healthcare). The images show a peak for each protein spot, with a peak height that is proportional to the spot intensity. (C: control; 1 h: Heat stress at 48°C for 1 h; 4 h: Heat stress at 48°C for 4 h)

**Table 1.**
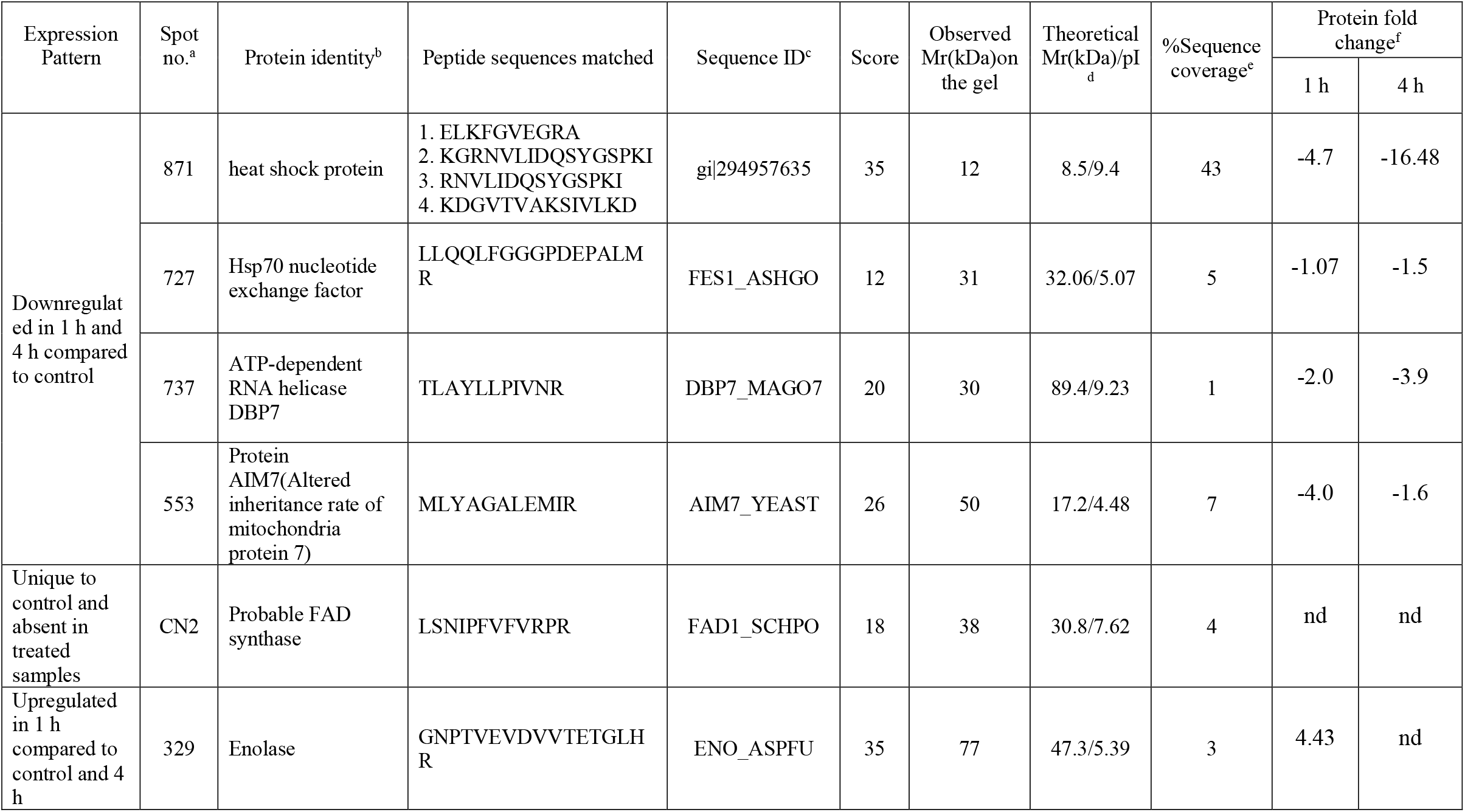
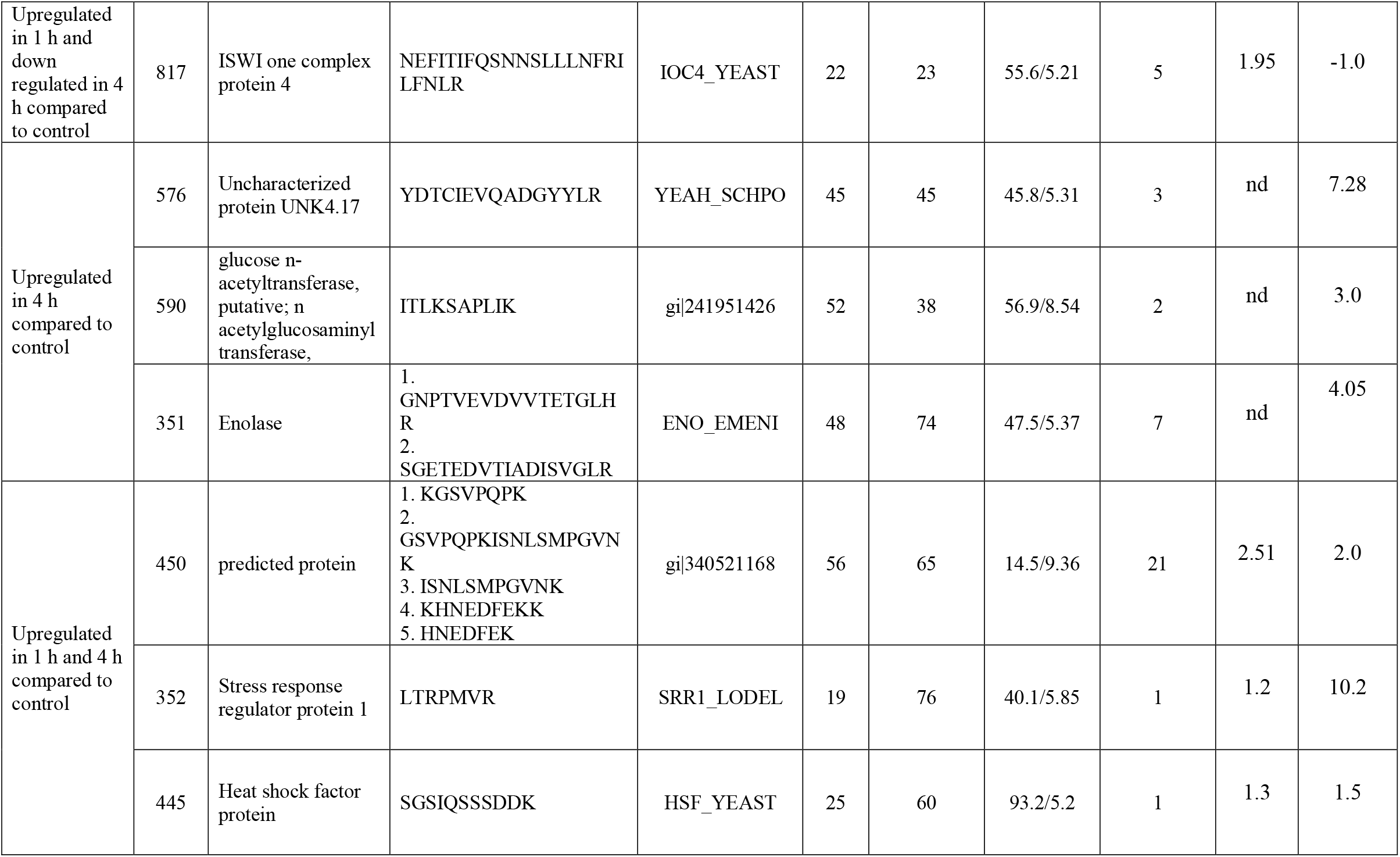
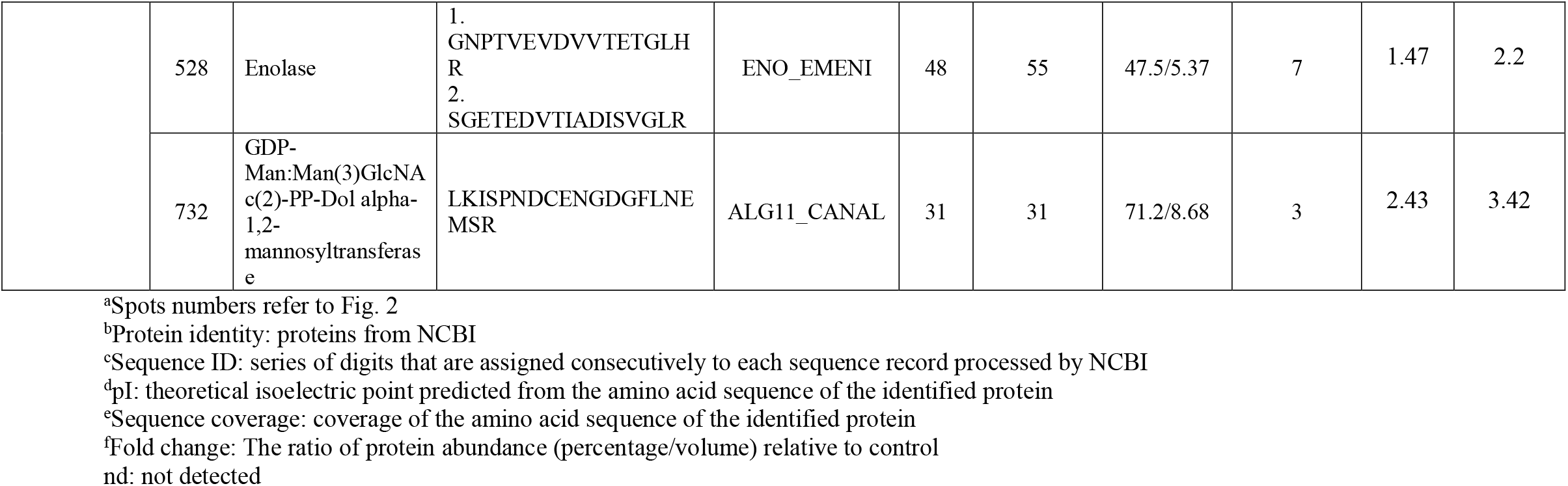
Differentially expressed proteins identified in *T. longibrachiatum* 673, TaDOR673 strain during heat stress through MALDI TOF-TOF

**Table 2.**
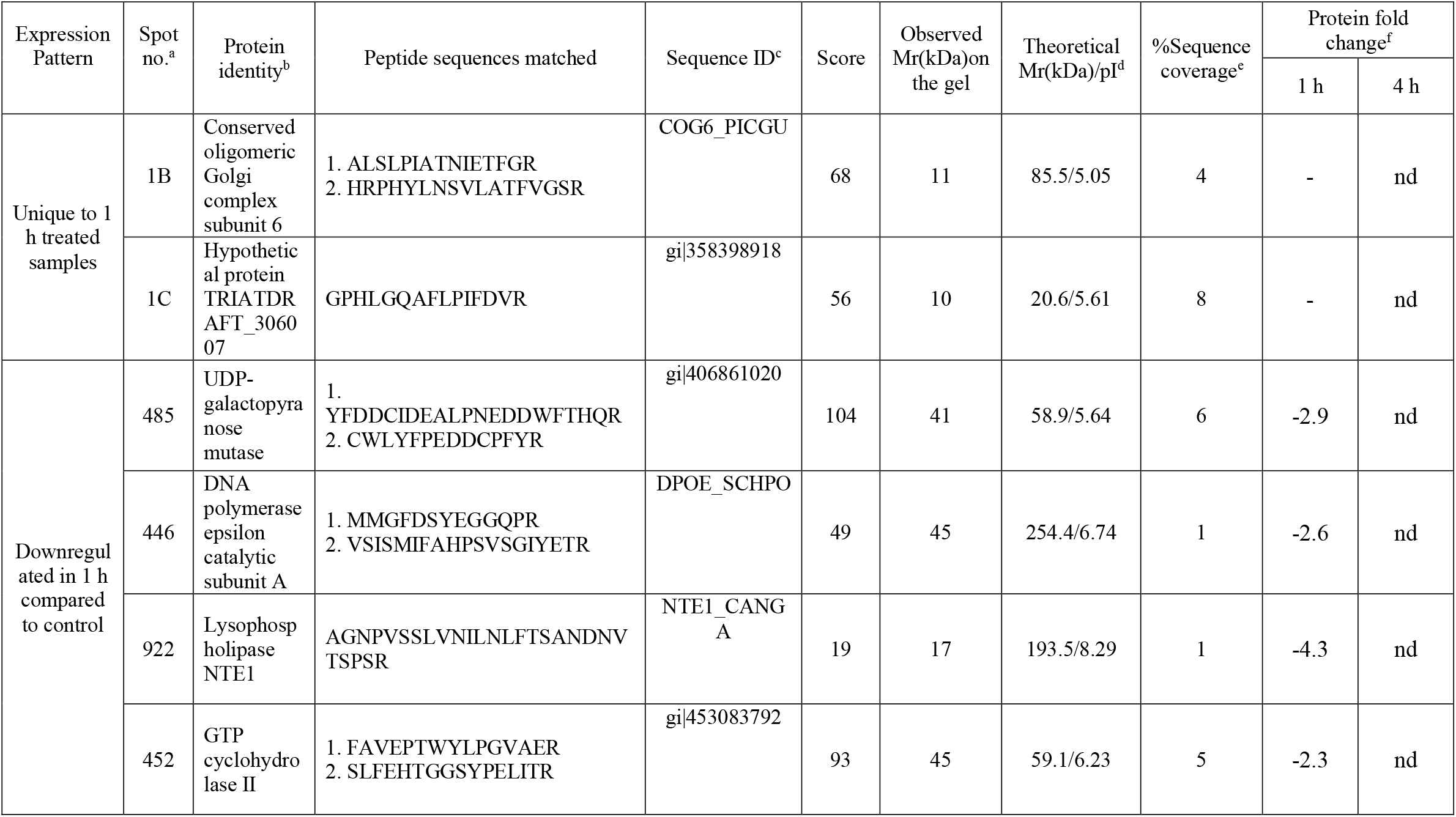
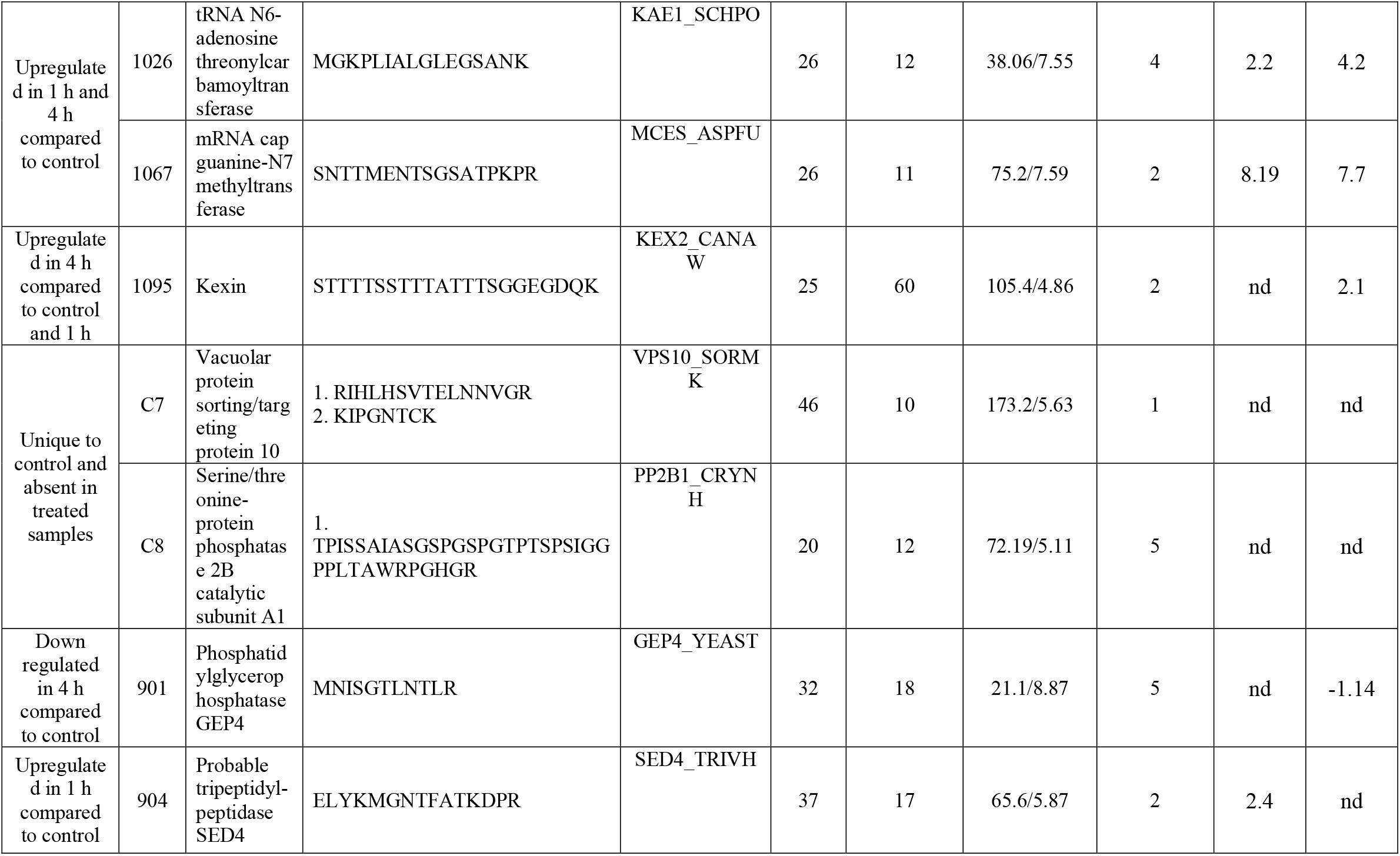
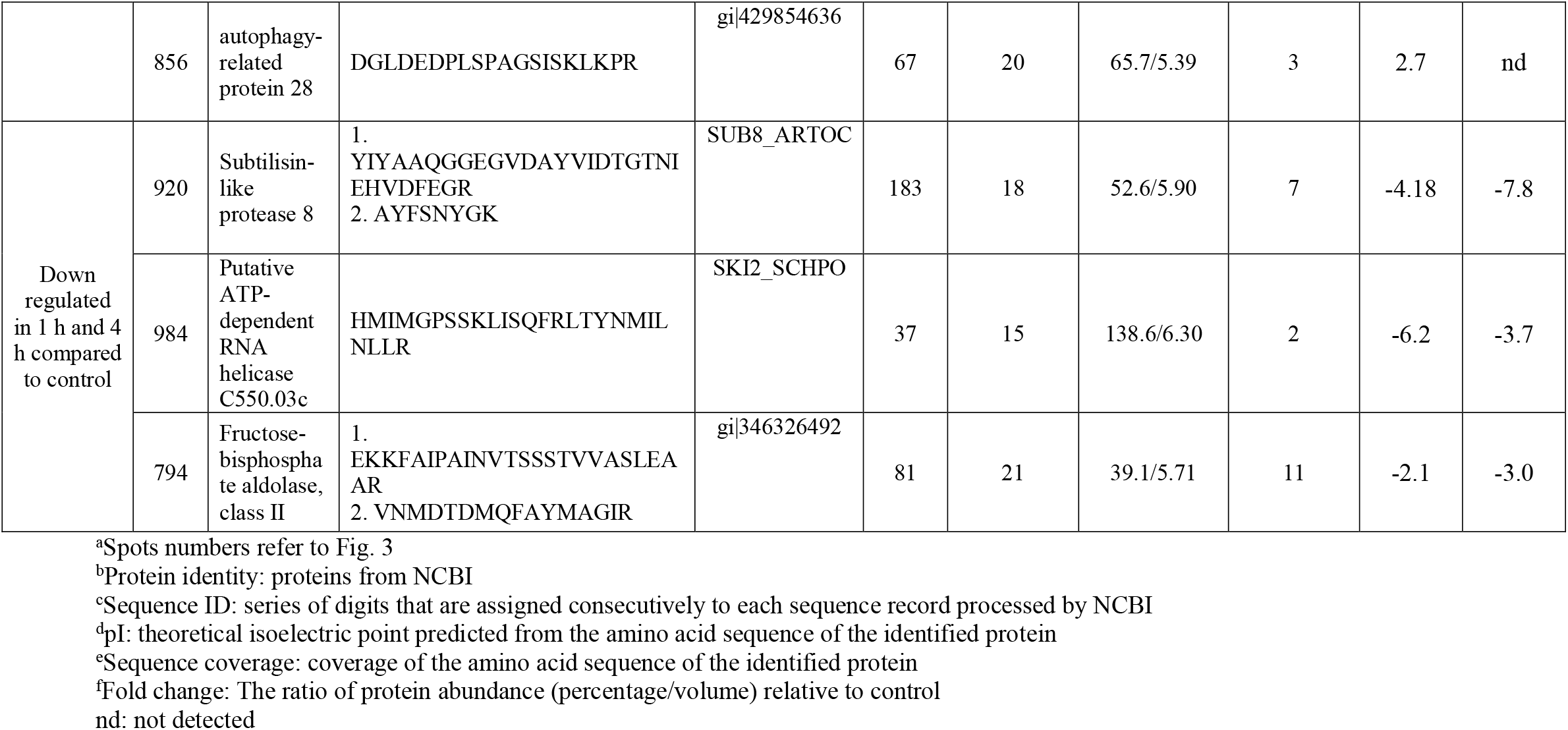
Differentially expressed proteins identified in *T. asperellum* 7316, TaDOR7316 proteins during heat stress through MALDI TOF-TOF

### 3.3. Differentially expressed proteins common in the two *Trichoderma* strains

#### 3.3.1. Proteins of cell wall re-modeling

There were differentially expressed proteins that were common between the two isolates tested (Table 1, 2 and supplementary table 2, 3). These proteins belonged to different cellular processes and are briefly discussed here.

Cell wall plays a crucial role in sensing and adapting to adverse stress conditions and earlier reports have shown that cell wall polysaccharides and lipid modifications contribute significantly to the induced heat and salt tolerance (Chatterjee et al. 2000). Composition of *Trichoderma* cell wall carbohydrates vary among species but are mainly composed of glucose, *N*-acetyl-glucosamine, *N*-acetylgalactosamine, galactose, and mannose (Prieto et al. 1997; Perlińska-Lenart et al. 2006). In TaDOR673, glucose N-acetyltransferase (Spot 590) which is involved in chitin synthesis was found to be absent after 1 h exposure but was up-regulated 3 fold after 4 h, indicating the activation of chitin biosynthesis during prolonged exposure to higher temperatures. Chitin constitutes 10-30% of fungal cell wall (de Nobel et al. 2000) and is actively synthesized during stress conditions (Georg & Gomes 2007; Ram et al. 2004). It has been hypothesized that N-acetylglucosamine (GlcNAc), a major component of fungal cell wall chitin might be a conserved cue to morphogenesis (Gilmore et al. 2013) and a mediator of cell signalling (Konopka 2012) during heat stress conditions.

Other cell wall proteins like α-1,2-mannosyltransferase (Spot 732) involved in cell wall biosynthesis was also found to be up-regulated 2 fold and 3 fold during 1 h and 4 h exposure, respectively, indicating that N-glycosylation, a major protein modification known to occur in different cells (Zhang et al. 2009), perhaps plays a crucial role in response to high temperature stress in this organism. However, in the moderately thermotolerant *Trichoderma* strain, TaDOR7316, a reduced expression of some of the proteins involved in cell wall re-modeling was observed. The enzyme, UDP-galactopyranose mutase (Spot 485) which is involved in the synthesis of the cell wall component, galactofuranose (Damveld et al. 2008), was down-regulated 3 fold after 1 h exposure and was completely absent at 4 h. Phospholipase B (PLB) protein (Spot 922) which is important for the turnover of phosphatidylcholine in cell membrane (Zaccheo et al. 2004) was also repressed (~ 4 fold) during high temperature stress. Physiological functions of fungal PLB are largely unknown but recently the role of a PLB homolog was identified as a mediator of osmotic stress response in fission yeast (Yang et al. 2003). The functional role of these proteins in imparting stress tolerance in TaDOR7316 strain needs to be empirically determined.

In TaDOR7316, membranes of cellular organelles were also impacted by high temperature stress. Phosphatidylglycerophosphatase GEP4 (Spot 901) that is involved in cardiolipin biosynthesis was completely absent during 1 h exposure to heat stress but was expressed at a low level (compared to control) during prolonged thermal stress. Cardiolipin levels seem to be important for osmoadaptation (Romantsov et al. 2009) and in cross-talk between mitochondria and vacuole (Chen et al. 2008). Thus in TaDOR7316 mitochondrial cell membrane signaling could be involved in activation of other stress response pathways that help it to ensure protection against thermal stress.

#### 3.3.2. Proteins of carbohydrate metabolism

Proteins of glycolysis and gluconeogenesis pathways are highly influenced during different stress conditions and the choice of the mechanism chosen by the organism to combat thermal stress mainly depends on the cytosol composition of the organism (Feofilova et al. 2000). This probably explains why different organisms choose either gluconeogenesis (Grossklaus et al. 2013; Kusch et al. 2007; Kim et al. 2007) or glucose catabolism (Schmidt et al. 2008; Barelle et al. 2006) to maintain cellular homeostasis during different stress conditions.

In TaDOR673, three isoforms of enolase (Spot 329, 528 and 351) were up regulated 2-4 fold during 1 h and 4 h exposure to heat stress (Table 1) and in agreement to our results, similar observations were made in several other fungi (George & Gomes 2007; Yin et al. 2009). Enolase being an important enzyme in carbohydrate metabolism and a major cell-envelope associated protein of *T. reesei* (Lim et al. 2001), we speculate that enolase could be involved in combating the heat stress conditions.

In contrast to above, in TaDOR7316 enzyme fructose 1, 6 bispohosphate aldolase (Spot 794) that catalyses a reversible reaction of conversion of fructose 1,6-bisphosphate to dihydroxyacetone phosphate and glyceraldehyde 3-phosphate was down regulated by 2 fold in 1 h and 3 fold in 4 h treated samples. Thus we hypothesize that this down regulation might lead to decreased use of the TCA cycle an increased glycerol biosynthesis and as observed in osmo-regulated cells of *Aspergillus nidulans* (Kim et al. 2007). This in turn would act as an osmoprotectant in protecting cells against dehydration or temperature stress (Feofilova et al. 2000).

#### 3.3.3. Proteins of Heat shock response (HSR) pathway

The role of *Trichoderma* derived heat shock proteins in stress tolerance has been well documented (Nicolas et al. 2014). and it is commonly observed that under various stress conditions, the heat shock proteins, Hsp70 and Hsp60 are coordinately expressed to ensure proper protein folding and maturation of nascent polypeptides (Kim et al. 2007; Yin et al. 2009).

Interestingly, in TaDOR673, Hsp70 nucleotide exchange factor Fes1 (Spot 727) and mitochondrial Hsp60 (Spot 871) were down regulated by 1.5 fold and 16 fold respectively during prolonged exposures to heat stress. The results were in agreement with earlier reports where they mentioned the reduced expression of Hsp60 in fungal cells exposed to pH stress (Schmidt et al. 2008). In normal cells, Fes1 specifically targets misfolded proteins recognized by Hsp70 in the ubiquitin/proteasome system (UPS) and cells lacking Fes1 are hypersensitive to induced protein mis-folding and display a strong and constitutive heat shock response mediated by heat shock transcription factor, *hsf1* (Abrams et al. 2014 & Gowda et al. 2013). Thus, induced expression of heat shock transcription factor (Spot 445) in TaDOR673 (~1.5 fold) probably indicates the activation of a constitutive heat shock response in the absence of Fes1 and we hypothesize that during high temperature stress, Hsp70 might use other dominant nucleotide exchange factors of the Hsp110 class to maintain the aggregation-prone proteins soluble in the cell, rather than targeting them to UPS for protein degradation. Moreover, it could also be possible that Hsf1 that was upregulated at higher temperatures might be involved in the diverse cellular processes like cell wall remodeling, carbohydrate metabolism, energy generation, etc during heat stress as suggested earlier (Imazu & Sakurai 2005; Yamamoto et al. 2005).

It was interesting to observe that small heat shock protein (sHsp), Hsp23 (Spot 1C) was induced during the initial hours of higher temperature in TaDOR7316 unlike in TaDOR673. Overexpression of Hsp23 correlated with increased thermotolerance in *T. harzianum* (Montero-Barrientos et al. 2007) reported earlier. In our study, overexpression of Hsp23 was confined only during the early hours of heat stress; we speculate that there could be an increased accumulation of denatured proteins in TaDOR7316 than TaDOR673 during the high temperature stress. These sHsps which are crucial in preventing stress-induced aggregation of partially denatured proteins could rescue this organism from heat stress and aid in restoration of the native conformations of denatured proteins when favorable conditions are restored. However, further studies are necessary to understand the role of these heat shock proteins in high temperature stress in these isolates.

#### 3.3.4. Proteins of cell signaling pathway

A hypothetical protein (Spot 352) in TaDOR673, which possessed a conserved domain for stress response regulator, was identified as an over-expressed protein at elevated temperatures (from 1 fold in 1 h to 10 fold in 4 h). This hypothetical protein could be the major stress signalling molecule of heat stress in TaDOR673 which in turn helps in the activation of other stress response genes. *Trichoderma* is known to use *Hog1*-mediated mitogen-activated protein kinase (MAPK) pathway in response to stress (Delgado-Jarana et al., 2006) in a two component phosphor-relay system where a phosphoryl group is transferred from a membrane bound sensor histidine kinase to an internal receiver domain. The phosphoryl group is then shuttled through a histidine phosphor-transfer protein to a terminal response regulator (RR). When stress signals such as oxidants, high salt, etc. are detected by cells, RR protein is dephosphorylated and is able to activate the downstream MAPK pathway to render cells adapt to the stress (Chauhan & Calderone 2008; Chauhan et al. 2006). Also, homologs of yeast RR are identified in other fungi and are found to be involved in stress adaptation (Desai et al. 2011). We therefore speculate that this hypothetical protein in TaDOR673 could probably be involved in the activation of MAPK pathway in response to high temperature.

However, in TaDOR7316 we observed the differential expression of calneurin which is a serine/threonine protein phosphatase. Calcineurin is activated by Ca^2+^-calmodulin by binding to the catalytic subunit, CnaA (Hashimoto et al., 1990). Interestingly in TaDOR7316 the catalytic subunit of calcineurin, which had a PP2B conserved domain (Spot C8) was completely absent during high temperature stress. But enough evidences are required to draw conclusions about the significance of these observations in high temperature tolerance.

#### 3.3.5. mRNA Stability

Other proteins that were common in the thermotolerant isolates were the RNA helicases. Two hypothetical proteins with conserved domains for helicases were found to be significantly downregulated in TaDOR673 (Spot 737) and TaDOR7316 (Spot 984) respectively. As is evident from Table 1 and 2, expression of RNA helicases was comparatively less in TaDOR7316 (~ 4-6 fold) than in TaDOR673 (~ 3 fold). The cellular requirement for increased RNA helicase abundance is generally associated with alteration of the stability of RNA secondary structure in response to the stress (Vinnemeier et al. 1999; Mukhopadhyay 2006) and our results corroborate other studies that showed that mutation in helicase function enhanced cellular tolerance to oxidative stress (Briolat & Reysset 2002).

In addition to maintaining mRNA structure, RNA helicases are also implicated in specialized processes including cell growth, differentiation, development, small RNA (sRNA) metabolism and response to abiotic stress (Rocak & Linder 2004; Owttrim 2006; Ambrus & Frolov 2009; Linder & Owttrim 2009). In this context, it is interesting that down regulation of RNA helicase expression in response to stress has rarely been reported (Owttrim 2006). Down regulation of RNA helicases in our study points to a mechanism that might be involved in induced stress tolerance when RNA helicases are either deleted/inactivated.

In spite of the reduced levels of RNA helicases, TaDOR7316 strain seem to protect its mRNA pool by increased expression (~8 fold) of other mRNA protecting enzymes (Spot 1067). This hypothetical protein probably identifies and hydrolyzes the aberrantly capped mRNA produced during heat stress in a manner similar to that observed in yeast (Jiao et al. 2010)

### 3.4. Proteins unique to TaDOR673

Apart from the major stress response pathways, proteins involved in actin depolymerization (Spot 553) and proteins of sulfur metabolism (Spot CN2) were downregulated during prolonged exposures to heat stress. As listed in Table 1, high temperature stress is probably ineffective on actin cytoskeleton during 1 h exposure but seems to be slowly affecting actin depolymerization at 4 h. Similar observations were made by Malerba et al. 2009 in yeast cells exposed to mild, moderate and high heat stress. We hypothesize that 48 °C is perhaps a moderate stress to TaDOR673 and therefore it probably did not undergo major cytoskeleton modifications at this level of heat stress.

A predicted protein with a conserved domain of PWWP (Spot 817) was upregulated in 1 h (~ 2 fold) but downregulated 1 fold after 4 h of heat stress. PWWP domain containing proteins possess DNA or histone binding activity (Yang and Everett, 2007) and to our knowledge this is the first report to describe the role of a chromatin remodeling protein in *Trichoderma* under high temperature stress. Chromatin remodeling is known to be essential to ensure the stress-inducible binding of transcription factors *viz*., Hsf1 to a majority of its targets (Hahn et al. 2004) and thus could act as a major regulator of the stress induced response of the fungus. This would be a good candidate gene for detailed analysis to understand the mechanism of heat stress tolerance.

In addition to these changes, increased expression (~7 fold) of oxidoreductases (Spot 576) was observed during prolonged heat stress conditions. As these enzymes mediate essential redox reactions in a cell, they may be important in response to several stresses (Grossklaus et al. 2013; Pusztahelyi et al. 2011). A hypothetical protein (Spot 450) was also identified to be deregulated during heat stress but their probable function could not be deciphered due to lack of conserved domain information (Supplementary table 2).

### 3.5. Proteins unique to TaDOR7316

#### 3.5.1. Unfolded protein response (UPR) and protein turnover

Among fungi, *Trichoderma* is predicted to contain a wealth of proteases (as predicted with use of the peptidase database MEROPS http://merops.sanger.ac.uk, Rawlings et al. 2012). Of these the dominant groups include aspartyl proteases, serine proteases, subtilisin-like proteases, dipeptidyl and tripeptidyl peptidases (Druzhinina et al. 2012). In the present study, two proteins belonging to protease S8 and S53 family were up regulated at higher temperature (Spot 1095; homologous to *T. virens* subtilisin like protease, EHK25893.1 and Spot 904; homologous to *T. reesei* tripeptidyl-peptidase 1 precursor, ETR98149.1). These proteins were up regulated ~2 fold both at 4 h and 1 h of heat treatments. Interestingly, a hypothetical protein (Spot 920), homologous to *T. atroviride* vacuolar serine protease, ABG57252.1 was found to be down regulated during heat stress. Therefore, in TaDOR7316, upregulation of these different proteases of UPR pathway could be essential to target the mis-folded or denatured proteins for degradation. Moreover, this process helps in both recycling of amino acids as well as clears the cellular matrix (Hilt 2004; Verghese et al. 2012).

Based on our results we speculate that under high temperature stress, misfolded proteins activate different enzymes of the unfolded protein response (UPR) pathway in the endoplasmic reticulum (ER) and in turn the UPR being an ER-to-nucleus signal transduction pathway might regulate a wide variety of target genes to maintain cellular homeostasis as reported earlier (Mori et al. 1996; Spear & Ng 2003).

#### 3.5.2. Vacuole biogenesis and Autophagy

We observed an increased expression of an autophagy protein, Apg6 (Spot 856) and two hypothetical proteins (Spot 1B and Spot 1026) with a conserved oligomeric-golgi domains (COG) during the stress conditions (Table 2). It was observed that under stress, when the ER-associated degradation (ERAD) pathway for misfolded proteins is saturable, cellular homeostasis is maintained by UPR pathway, which transports the excess substrate (misfolded proteins) to vacuole for protein turnover (Spear & Ng 2003). Studies have shown that endocytic pathways, proteosome pathways and regulation of autophagy are induced during heat stress in filamentous biocontrol fungi (Wang et al. 2014). COG complex which helps in the formation of double-membrane sequestering vesicles during autophagy (Yen et al. 2010) is principally important for retrograde trafficking within the Golgi complex and possibly for ER transport to Golgi and endosome transport to Golgi complex (Whyte & Munro 2001; Ram et al. 2002; Bruinsma et al. 2004; Oka et al. 2004; Zolov & Lupashin 2005).

Thus we hypothesized that under heat stress TaDOR7316 probably tries to maintain the native conformation of proteins during early hours of exposure by inducing the expression of heat shock proteins (Hsp23 as mentioned in above sections) but during prolonged exposures to heat stress, it probably maintains cellular homeostasis by deploying UPR pathway to transport the increased levels of denatured/misfolded proteins to vacuoles for degradation.

Several vacuolar protein sorting complexes exist in cells to help efficient retrograde transport of proteins from endosome-to-Golgi complex. Their role in stress resistance, host cell interactions, and virulence are also well documented (Liu et al. 2014). In the present study, expression of a vacuolar sorting protein with a conserved domain of VPS10 was identified in unstressed samples but was completely absent under stress conditions in TaDOR7316 (Spot C7). The VPS10 encodes a Carboxypeptidase Y (CPY) sorting receptor that executes multiple rounds of sorting by cycling between the late Golgi and a pre-vacuolar endosome-like compartment. Studies have shown that mutations in VPS10 resulted in selective mis-sorting and secretion of CPY but had no impact on the delivery of other vacuolar proteins (Marcusson et al. 1994). Thus we speculate that in TaDOR7316, in spite of the absence of certain vacuolar sorting signal molecules, the fate of most of the misfolded proteins might be taken care of by the proteins of UPR pathway and that in turn might trigger vacuole biogenesis and autophagy to maintain protein turnover and cellular homeostasis.

### 3.6. Other proteins

Protein involved in DNA replication (spot 446, homologous to *T. reesei* QM6a DNA polymerase epsilon, XP_006964819.1) was downregulated by 2 fold in 1 h and completely repressed during 4 h exposure to heat stress in TaDOR7316. Although little is known about its effects on DNA integrity and DNA replication process, it has been shown that in yeast the absence of topoisomerase activities did not prevent induction of either heat or radiation resistance. However, if both topoisomerase I and II activities were absent, sensitivity of yeast to become thermally tolerant was markedly increased (Boreham et al. 1990).

We identified a hypothetical protein (Spot 452) homologous to GTP cyclohydrolase of *T. reesei* QM6a whose expression was repressed by 2 fold during 1 h and was completely absent at 4 h of heat stress. GTP cyclohydrolase is involved in riboflavin biosynthesis and in turn riboflavin protects cells from stress injuries (Perumal et al. 2005; Sugiyama 1991) Elevated riboflavin is required for post-photoinductive events in sporulation of a *Trichoderma* auxotroph (Horwitz & Gressel 1983) and down regulation of this enzyme probably explains the reduced sporulation observed in TaDOR7316 during the heat stress. Similar results were also observed in *A. nidulans* and *Ashbya gossypii* exposed to oxidative stress agents (Pusztahelyi et al. 2011; Kavitha & Chandra 2009). However, further studies are required to understand the exact role of these proteins in heat stress tolerance.

To gain further insight into the mechanism of thermotolerance in the two strains used in this study, we have also carried out microarray analysis to profile the altered expression in transcripts due to higher temperature (Poosapati et al. Manuscript under preparation). By comparing the proteomic and transcriptome data, we would like to establish the correlation between transcript and protein levels.

## 4. Conclusion

In our study, we compared proteins profiles of two different thermotolerant isolates of *Trichoderma* under heat stress. These isolates differ in the level of thermotolerance and as hypothesized we identified some putative heat stress responsive proteins that are probably responsible for the thermotolerance levels of TaDOR673 and TaDOR7316. We propose that in TaDOR673, under heat stress, the organism undergoes cell wall changes and metabolic changes like increased chitin production and increased enolase and trehaolse biosynthesis. In addition, stress signals were sensed by putative cell membrane sensors which in turn transfer the signals to response regulators (RR) and activate MAPK signalling pathway. HSR pathway also seems to play a crucial role in the thermotolerance of TaDOR673. Based on the proteomic changes, we hypothesize that Hsf1 might be involved in cell wall remodeling to combat the heat stress in TaDOR673 strain. However, in the moderately thermotolerant isolate TaDOR7316, heat stress probably resulted in increased accumulation of misfolded proteins and in order to protect the cell from toxicity, misfolded proteins were targeted to degradation through UPR and autophagy. Moreover, sHsps like Hsp23, were found to be a part of early stress responses in TaDOR7316.

Thus, we observed that the superior thermotolerant isolate, TaDOR673 was able to manage the levels of heat stress through the activation of stress signaling and heat shock response pathways whereas the moderately thermotolerant isolate, TaDOR7316 experienced severe stress during prolonged exposures to higher temperatures and the strain circumvents the heat stress by getting rid of toxic proteins to maintain cellular homeostasis. Further studies are required to empirically establish the functional role of the differentially expressed proteins in these two strains during exposure to higher temperatures.

## Acknowledgement

The present research work was carried out with the research grant (Project no.C2082) given by National Agricultural Innovation Program, ICAR, New Delhi, India to RDP and VDK as well as the Fellowship to SP. Facilities extended by the Director, Indian Institute of Oilseeds Research (formerly Directorate of Oilseeds Research), Rajendranagar, Hyderabad, India for carrying out this research is acknowledged. The authors would also like to thank Dr. Lekha Dinesh Kumar, CCMB, Habsiguda, Mr. Velu Mani Selvaraj and Mr. Anil Kumar, Research Scholars at IIOR, Hyderabad for their technical support.

