## Supplementary Figure 1. for "Proteomic analysis reveals different sets of proteins expressed during high temperature stress in two thermotolerant isolates of *Trichoderma*"

### Slide 1
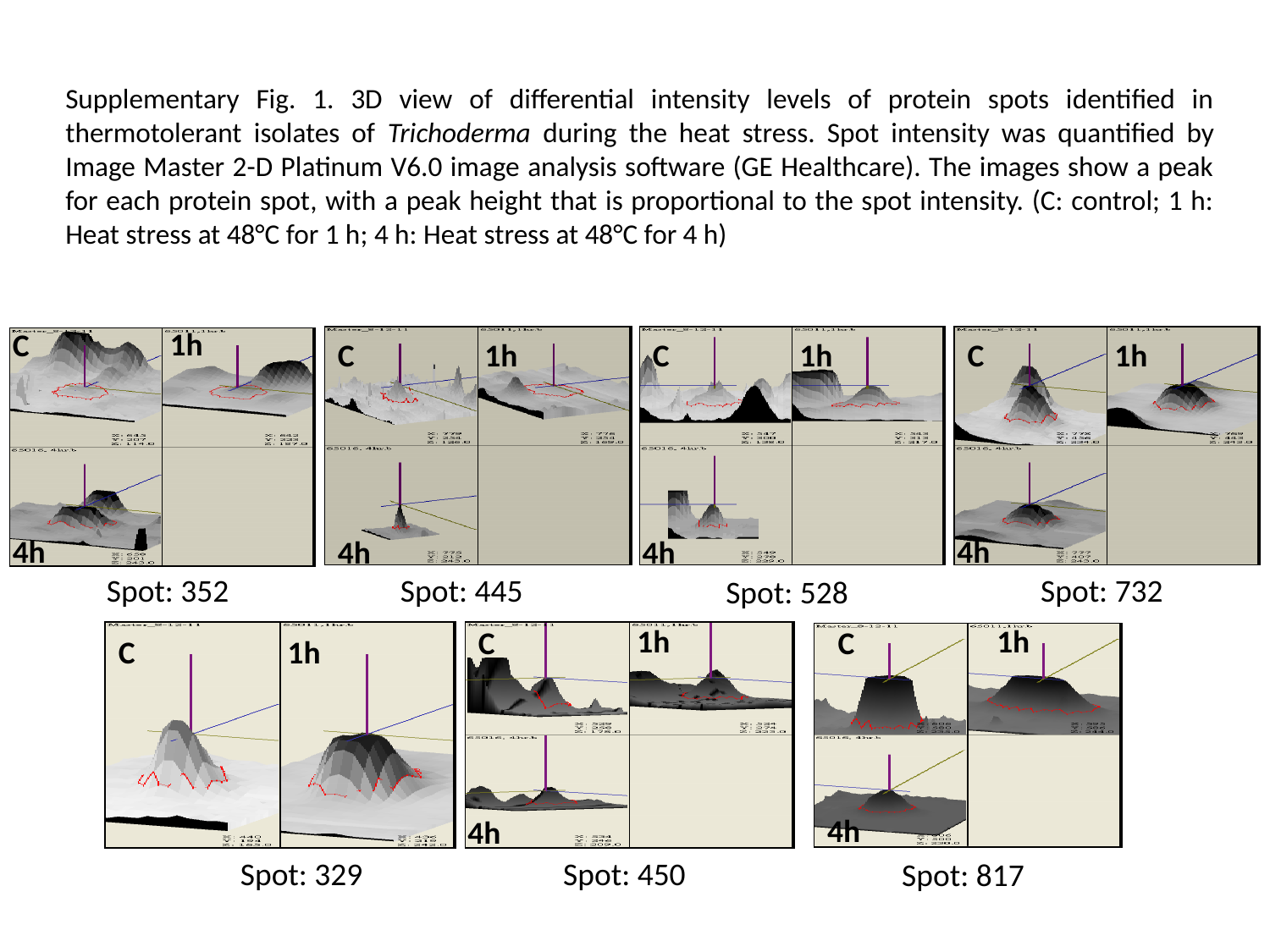

Supplementary Fig. 1. 3D view of differential intensity levels of protein spots identified in thermotolerant isolates of Trichoderma during the heat stress. Spot intensity was quantified by Image Master 2-D Platinum V6.0 image analysis software (GE Healthcare). The images show a peak for each protein spot, with a peak height that is proportional to the spot intensity. (C: control; 1 h: Heat stress at 48°C for 1 h; 4 h: Heat stress at 48°C for 4 h)
1h
C
Spot: 352
Spot: 445
Spot: 732
Spot: 528
C
1h
C
1h
C
1h
4h
4h
4h
4h
1h
C
Spot: 329
Spot: 450
C
1h
4h
1h
C
Spot: 817
4h

### Slide 2
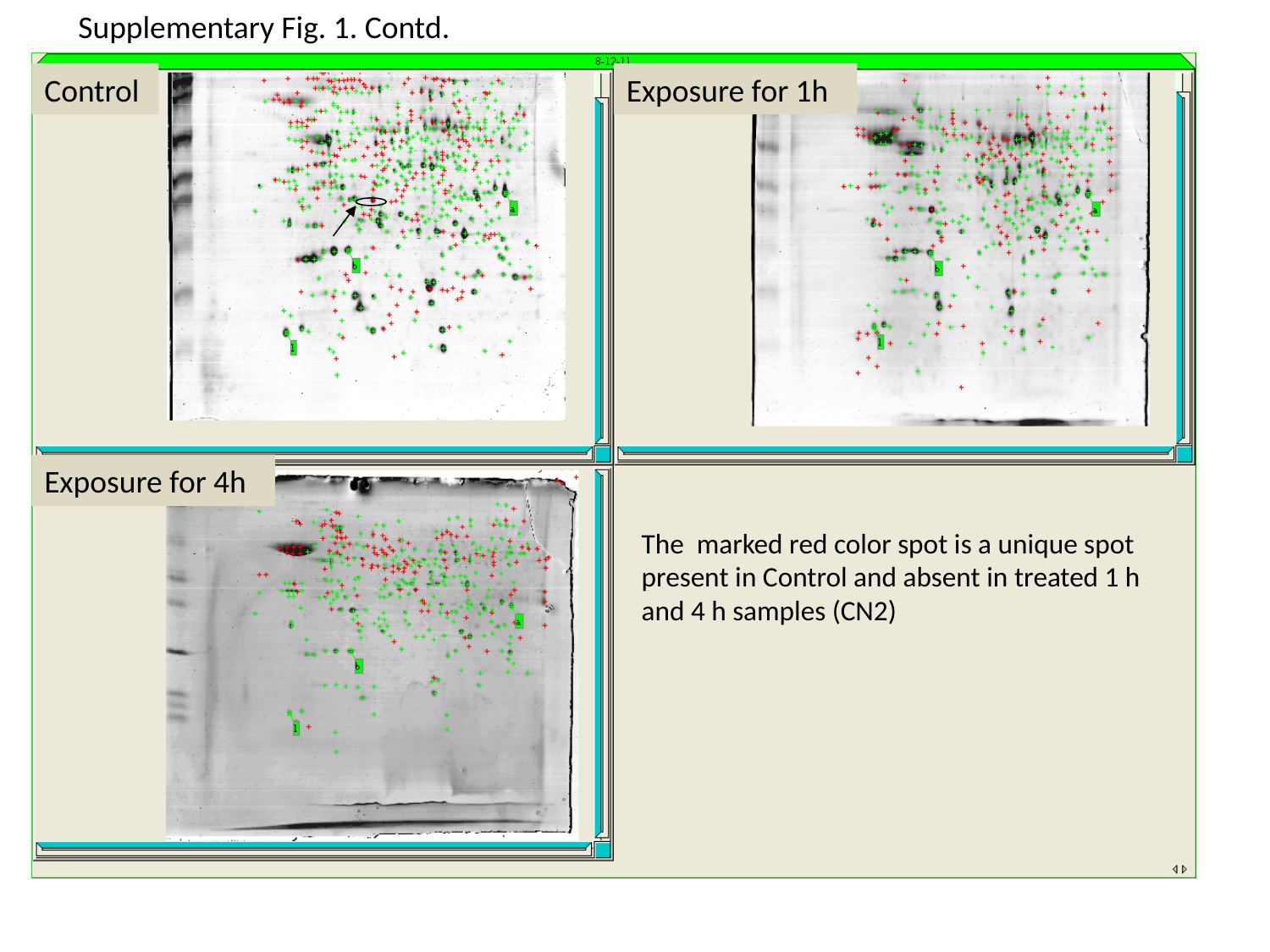

Supplementary Fig. 1. Contd.
The marked red color spot is a unique spot present in Control and absent in treated 1 h and 4 h samples (CN2)
Control
Exposure for 1h
Exposure for 4h

### Slide 3
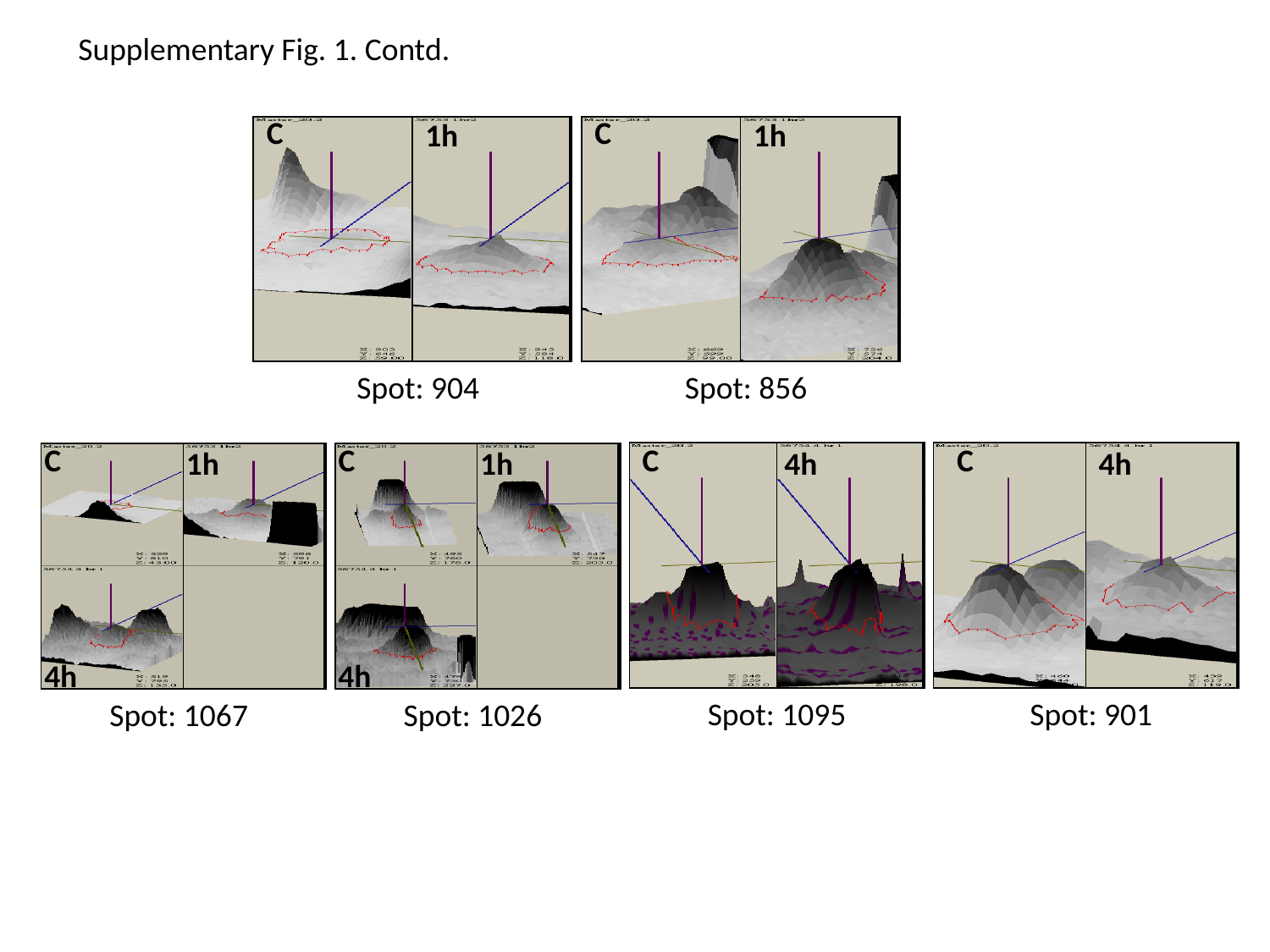

Supplementary Fig. 1. Contd.
C
C
1h
1h
Spot: 904
Spot: 856
C
C
1h
1h
Spot: 1067
Spot: 1026
4h
4h
C
4h
Spot: 1095
C
4h
Spot: 901

### Slide 4
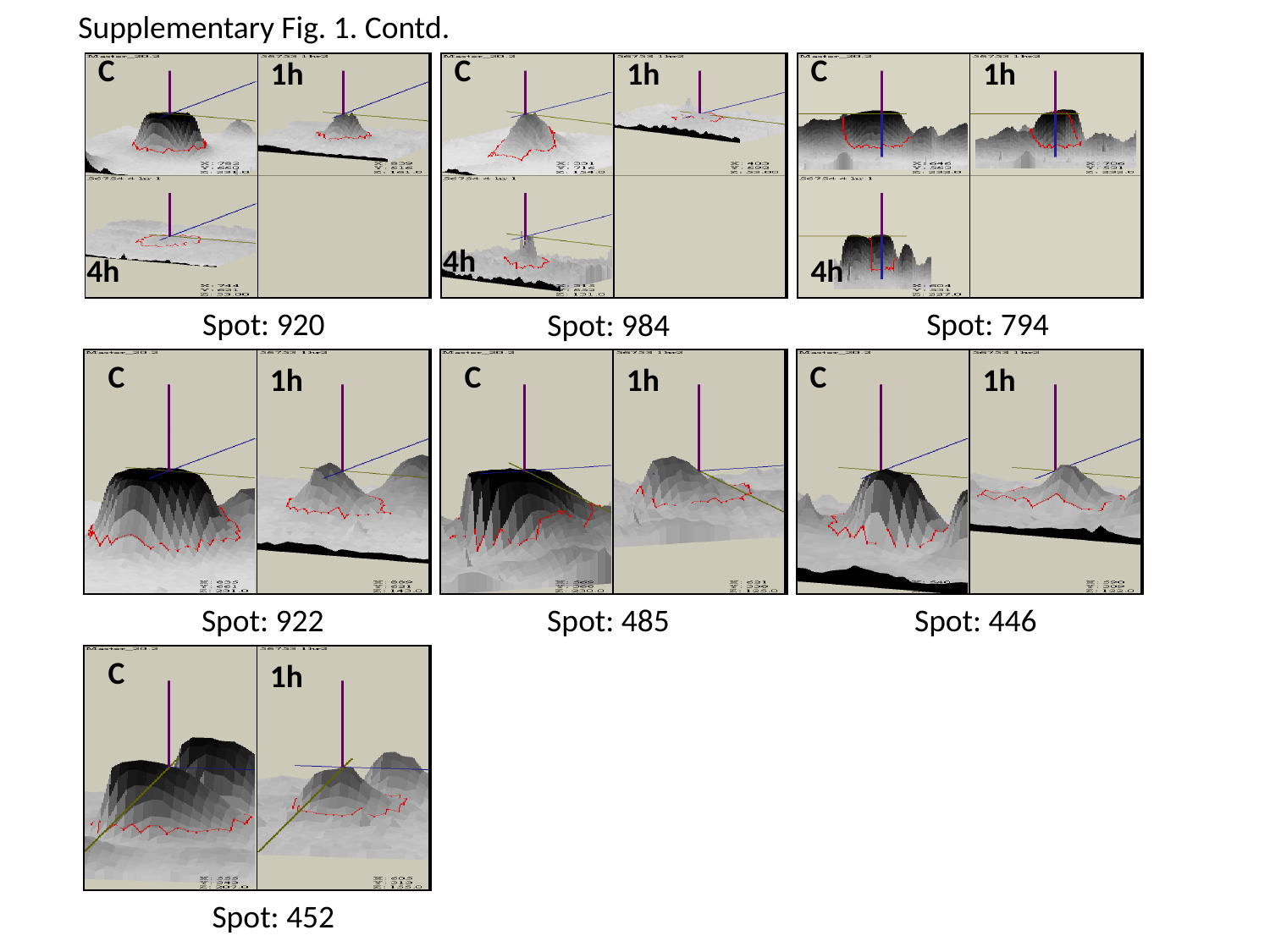

Supplementary Fig. 1. Contd.
C
C
C
1h
1h
1h
Spot: 920
Spot: 794
Spot: 984
4h
4h
4h
Spot: 922
Spot: 485
Spot: 446
Spot: 452
C
C
C
1h
1h
1h
C
1h

### Slide 5
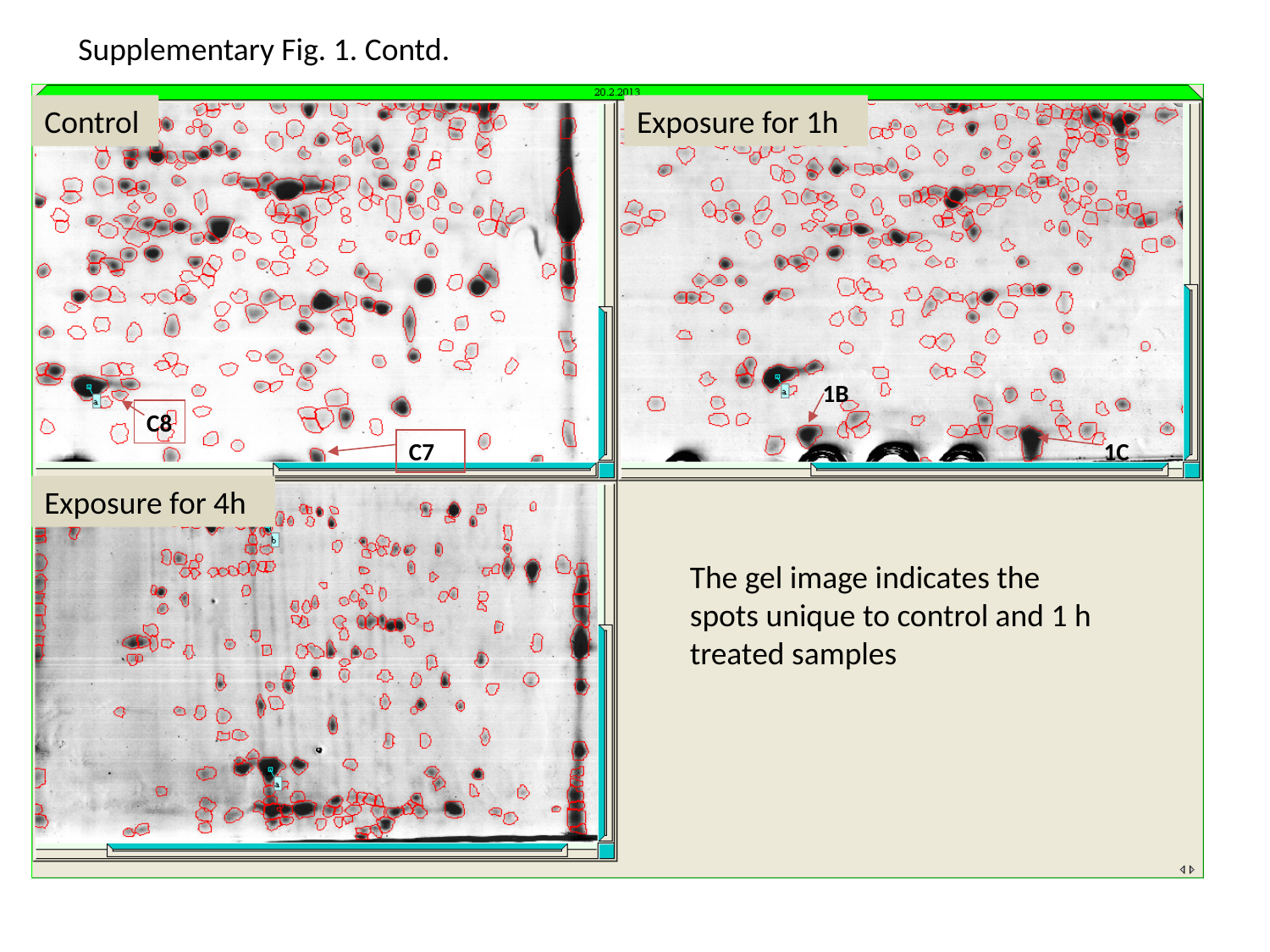

Supplementary Fig. 1. Contd.
1B
C8
C7
1C
Control
Exposure for 1h
Exposure for 4h
The gel image indicates the spots unique to control and 1 h treated samples
