## Supplementary Table 1 for "Proteomic analysis reveals different sets of proteins expressed during high temperature stress in two thermotolerant isolates of *Trichoderma*"

**Supplementary table 1.** List of accession numbers of thermotolerant *Trichoderma* isolates

| **Isolate** | **Accession numbers** | |
| --- | --- | --- |
| Gene | TEF1α | RNA Polymerase subunit B |
| *T. asperellum* 7316, TaDOR7316 | KM190858 | KX389492 |
| *T. longibrachiatum* 673, TaDOR673 | KM190859 | KX389493 |
