## Supplementary Table 2 & 3 for "Proteomic analysis reveals different sets of proteins expressed during high temperature stress in two thermotolerant isolates of *Trichoderma*"

Supplementary table 2. BLASTP and CDD (Conserved domain) analyses of protein identified during heat stress in *T. longibrachiatum* 673, TaDOR673

| Expression Pattern | Spot  no. | NCBI Hit of BlastP against *Trichoderma* | Fold-regulation | | Conserved domains | Accession No. | Score | E-value | %Identity |
| --- | --- | --- | --- | --- | --- | --- | --- | --- | --- |
|  |  |  | 1 h | 4 h |  |  |  |  |  |
| Downregulated in 1 h and 4 h compared to control | 871 | hsp60 mitochondrial precursor-like protein [*Trichoderma* *reesei* QM6a] | -4.7 | -16.48 | GroEL domain | [XP_006961679.1](http://www.ncbi.nlm.nih.gov/protein/589099313?report=genbank&log$=prottop&blast_rank=2&RID=JU43BRPF013) | 140 | 2e-40 | 86% |
|  | 727 | hypothetical protein TRIVIDRAFT_185457 [*Trichoderma* *virens* Gv29-8] | -1.07 | -1.5 | FES1 and ARM domain | EHK27263.1 | 143 | 2e-40 | 44% |
|  | 737 | predicted protein [*Trichoderma* *reesei* QM6a] | -2.0 | -3.9 | Helicase domains | XP_006961979.1 | 716 | 0.0 | 49% |
|  | 553 | glia maturation factor beta [*Trichoderma* *reesei* RUT C-30] | -4.0 | -1.6 | Actin depolymerization factor/cofilin-like domains (ADF domains) | ETS05399.1 | 95.9 | 1e-26 | 39% |
| Unique to control and absent in treated samples | CN2 | hypothetical protein TRIATDRAFT_38099 [*Trichoderma* *atroviride* IMI 206040] | nd | nd | Phosphoadenosine phosphosulfate reductase family | EHK45271.1 | 184 | 4e-56 | 42% |
| Upregulated in 1 h compared to control and 4 h | 329 | phosphopyruvate hydratase [*Trichoderma* *reesei* QM6a] | 4.43 | nd | Enolase | XP_006963341.1 | 795 | 0.0 | 88% |
| Upregulated in 1 h and down regulated in 4 h compared to control | 817 | predicted protein [*Trichoderma* *reesei* QM6a] | 1.95 | -1.0 | PWWP domain | XP_006966270.1 | 51.6 | 1e-06 | 26% |
| Upregulated in 4 h compared to control | 576 | hypothetical protein TRIVIDRAFT_181102 [*Trichoderma* *virens* Gv29-8] | nd | 7.28 | Oxidoreductase family | EHK20255.1 | 31.2 | 1.8 | 24% |
|  | 590 | glycosyltransferase family 8 protein [*Trichoderma* *atroviride* IMI 206040] | nd | 3.0 | Glucosyl transferase 8 (GT8) family | EHK46197.1 | 105 | 1e-24 | 23% |
|  | 351 | phosphopyruvate hydratase [*Trichoderma* *reesei* QM6a] | nd | 4.05 | Enolase superfamily | EHK50091.1 | 790 | 0.0 | 87% |
| Upregulated in 1 h and 4 h compared to control | 450 | predicted protein[*Trichoderma* *reesei* QM6a] | 2.51 | 2.0 | No conserved domains | XP_006962894.1 | 260 | 1e-86 | 100% |
|  | 352 | hypothetical protein TRIVIDRAFT_30731 [*Trichoderma* *virens* Gv29-8] | 1.2 | 10.2 | Signal receiver domain | EHK24432.1 | 59.7 | 2e-09 | 29% |
|  | 445 | heat shock factor-type DNA-binding domain-containing protein [*Trichoderma* *reesei* QM6a] | 1.3 | 1.5 | Heat shock transcription factor | XP_006965998.1 | 95.9 | 5e-20 | 27% |
|  | 528 | Phosphopyruvate hydratase [*Trichoderma* *reesei* QM6a] | 1.47 | 2.2 | Enolase | XP_006963341.1 | 795 | 0.0 | 88% |
|  | 732 | Glycosyltransferase family 4 protein [*Trichoderma* *atroviride* IMI 206040] | 2.43 | 3.42 | Glycosyltransferase group 1 family | EHK45433.1 | 233 | 4E-68 | 34% |

nd: Not detected

1 h: Heat stress treatment at 48°C for 1 h

4 h: Heat stress treatment at 48°C for 4 h

Supplementary table 3. BLASTP and CDD (Conserved domain) analyses of protein identified during heat stress in *T. asperellum* 7316, TaDOR7316

| Expression Pattern | Spot  no. | NCBI Hit of BlastP of protein against *Trichoderma* | Fold-regulation | | Conserved domains | Accession No. | Score | E-value | %Identity |
| --- | --- | --- | --- | --- | --- | --- | --- | --- | --- |
|  |  |  | 1 h | 4 h |  |  |  |  |  |
| Unique to 1 h treated samples | 1B | Intra-Golgi transport complex, subunit 6 [*Trichoderma* *reesei* QM6a] | - | nd | Conserved oligomeric complex (COG6) | XP_006965592.1 | 99.8 | 2e-21 | 21% |
|  | 1C | small heat shock protein [*Trichoderma* *harzianum*] | - | nd | Hsp20/alpha crystalline family | AAX55622.1 | 97.1 | 2e-24 | 37% |
| Downregulated in 1 h compared to control | 485 | hypothetical protein TRIATDRAFT_301560 [*Trichoderma* *atroviride* IMI 206040] | -2.9 | nd | NAD(P)-binding Rossmann-like domain | EHK40773.1 | 855 | 0.0 | 77% |
|  | 446 | DNA polymerase epsilon, catalytic subunit A [Trichoderma reesei QM6a] | -2.6 | nd | DNA polymerase type-B epsilon subfamily catalytic domain | XP_006964819.1 | 2130 | 0.0 | 48% |
|  | 922 | hypothetical protein TRIATDRAFT_87546 [Trichoderma atroviride IMI 206040] | -4.3 | nd | No conserved domains | EHK44512.1 | 22.3 | 4.8 | 86% |
|  | 452 | GTP cyclohydrolase [*Trichoderma* *reesei* QM6a] | -2.3 | nd | GTP cyclohydrolase II (RibA) | XP_006967668.1 | 728 | 0.0 | 68% |
| Upregulated in 1 h and 4 h compared to control | 1026 | hypothetical protein TRIATDRAFT_52866 [*Trichoderma* *atroviride* IMI 206040] | 2.2 | 4.2 | tRNA threonylcarbamoyl adenosine modification protein YeaZ; Inactive homolog of metal-dependent proteases (COG1214); Glycoprotease family (Peptidase_M22) | EHK50571.1 | 498 | 2e-176 | 69% |
|  | 1067 | hypothetical protein TRIATDRAFT_265826 [*Trichoderma* *atroviride* IMI 206040] | 8.19 | 7.7 | mRNA capping enzyme (pfam03291); SAM-dependent methyltransferases | EHK44240.1 | 494 | 4e-168 | 58% |
| Upregulated in 4 h compared to control and 1 h | 1095 | subtilisin like protease [*Trichoderma* *virens* Gv29-8] | nd | 2.1 | Peptidase S8 family domain in Protein convertases; | EHK25893.1 | 564 | 0.0 | 43% |
| Unique to control and absent in treated samples | C7 | vacuolar sorting protein [*Trichoderma* *reesei* QM6a] | nd | nd | Non-viral sialidases; VPS10 domain | XP_006962518.1 | 1559 | 0.0 | 53% |
|  | C8 | Serine/threonine-protein phosphatase 2B catalytic subunit [*Trichoderma* *reesei* RUT C-30] | nd | nd | PP2B, metallophosphatase domain | ETS03163.1 | 828 | 0.0 | 78% |
| Down regulated in 4 h compared to control | 901 | predicted protein [*Trichoderma* *reesei* QM6a] | nd | -1.14 | Mitochondrial PGP phosphatase | XP_006964630.1 | 181 | 2e-57 | 50% |
| Upregulated in 1 h compared to control | 904 | tripeptidyl-peptidase 1 precursor [*Trichoderma* *reesei* RUT C-30] | 2.4 | nd | Peptidase domain in the S53 family | ETR98149.1 | 377 | 4e-122 | 41% |
|  | 856 | hypothetical protein TRIVIDRAFT_213745 [*Trichoderma* *virens* Gv29-8] | 2.7 | nd | chromosome segregation protein SMC; Autophagy protein Apg6 | EHK19294.1 | 434 | 1e-143 | 45% |
| Down regulated in 1 h and 4 h compared to control | 920 | vacuolar serine protease [*Trichoderma* *atroviride*] | -4.18 | -7.8 | Peptidase S8 family domain | ABG57252.1 | 642 | 0.0 | 65% |
|  | 984 | hypothetical protein TRIVIDRAFT_209259 [*Trichoderma* *virens* Gv29-8]; antiviral helicase [*Trichoderma* *reesei* RUT C-30] | -6.2 | -3.7 | DEAD-like helicases superfamily | EHK22776.1; ETS02258.1 | 1116; 427 | 0.0;  8e-130 | 49%;  41% |
|  | 794 | fructose bisphosphate aldolase [*Trichoderma* *reesei* QM6a] | -2.1 | -3.0 | Class II Type A, Fructose-1,6-bisphosphate (FBP) aldolases | XP_006968688.1 | 661 | 0.0 | 91% |

nd: Not detected

1 h: Heat stress treatment at 48°C for 1 h

4 h: Heat stress treatment at 48°C for 4 h
